# Moving beyond neurons: the role of cell type-specific gene regulation in Parkinson’s disease heritability

**DOI:** 10.1101/442152

**Authors:** Regina H Reynolds, Juan Botía, Mike A Nalls, International Parkinson’s Disease Genomics Consortium (IPDGC), System Genomics of Parkinson’s Disease (SGPD), John Hardy, Sarah A Gagliano, Mina Ryten

## Abstract

Parkinson’s disease (PD), with its characteristic loss of nigrostriatal dopaminergic neurons and deposition of α-synuclein in neurons, is often considered a neuronal disorder. However, in recent years substantial evidence has emerged to implicate glial cell types, such as astrocytes and microglia. In this study, we used stratified LD score regression and expression-weighted cell-type enrichment together with several brain-related and cell-type-specific genomic annotations to connect human genomic PD findings to specific brain cell types. We found that PD heritability does not enrich in global and regional brain annotations or brain-related cell-type-specific annotations. Likewise, we found no enrichment of PD susceptibility genes in brain-related cell types. In contrast, we demonstrated a significant enrichment of PD heritability in a curated lysosomal gene set specifically expressed in astrocytic and microglial subtypes. Our results suggest that PD risk loci do not lie in specific cell types or individual brain regions, but rather in global cellular processes to which cell types may have varying vulnerability.

---

Late-onset sporadic forms of neurodegenerative diseases are devastating conditions imposing an increasing burden on healthcare systems worldwide. Currently, 2-3% of the population over 65 years of age are living with Parkinson’s disease (PD), making this disorder the most prevalent late-onset neurodegenerative disorder worldwide after Alzheimer’s disease^1^. This progressive condition is characterised by the loss of dopaminergic neurons in the substantia nigra pars compacta manifesting clinically as a tremor at rest, muscle rigidity and bradykinesia^1,2^. Existing symptomatic treatments do not alter the course of the disease and their effectiveness declines with time, which makes the identification of potential therapeutic targets of key importance.

The primary focus of PD research to date has been on neurons and, more specifically, nigrostriatal dopaminergic neurons. This focus is driven in part because the death of dopaminergic neurons is primarily responsible for the motor features of PD, but also because the most prominent and distinctive neuropathological findings in PD are the presence of neuronal inclusions, termed Lewy bodies^1,2^. The findings that alpha synuclein (encoded by the gene *SNCA*) is predominantly expressed in neurons^2,3^, is the major component of Lewy bodies^3,4^, and mutations in *SNCA* give rise to autosomal dominant PD^5–8^ provide a key link between *SNCA* function, neurons and disease pathogenesis. Furthermore, the identification of risk SNPs at the *SNCA* locus through genome-wide association studies (GWAS) of sporadic PD^9^ provides support for the importance of *SNCA*-related pathways and, by implication, neurons in both sporadic and Mendelian forms of PD. Despite this neuronal focus, there is also growing evidence to suggest the involvement of other cell types in PD pathogenesis. In particular, astrocytes and microglia have been highlighted^10,11^; for instance, with a recent study demonstrating that blocking the microglial-mediated conversion of astrocytes to an A1 neurotoxic phenotype was neuroprotective in mouse models of sporadic and familial α-synucleinopathy^12^.

In previous work, we applied stratified LD score regression and gene-set enrichment methods to determine if particular functional marks for regulatory activity and gene-set lists were enriched for sporadic PD genetic heritability^13^. We did not observe enrichment for the various brain annotations assessed (this did not include brain-relevant cell types) and in fact found further evidence for the importance of the adaptive and innate immune system.

The increasing power of GWASs (with the most recently published PD GWAS including 20,184 cases and 397,324 controls, resulting in over 35 associated loci^9^) coupled with the increased availability of cell-specific gene expression data provides a new opportunity to address the potential cellular-specificity of disease heritability, as was elegantly demonstrated for schizophrenia in a study by Skene et al.^14^. Brain regions contain a mixture of cell types, such as neurons, microglia and astrocytes, which may exhibit their own specific regulatory features that could be masked when averaging features across cell types. Resolving this question has become increasingly important; with the advent of induced pluripotent stem cell models of disease, modelling PD *in vitro* is now possible, and this implies some decision about the cell type of interest. In this study, we addressed cellular heterogeneity through the analysis of genomic regions overlapping regulatory marks or gene expression from cell types within the brain, including neurons. We focus on PD GWAS datasets and use schizophrenia (SCZ) GWAS datasets for comparisons purposes.

## Results

To study the cellular specificity of the heritability of sporadic PD, we compiled brain-related genomic annotations denoting tissue- and cell-type-specific markers of activity. We used several approaches to capture the expression profiles of human brain-related cell types (Figure 1, see Methods). This was because no single data set had all the desirable properties; namely, data that was human in origin, covered multiple brain regions, had high cellular detail and was derived from large numbers of individuals. Using the largest publicly available GWASs of PD^15^ and SCZ^16^, we applied stratified LD score regression (LDSC) to assess enrichment of the common SNP heritability of PD and SCZ, respectively, for each annotation category. SCZ heritability has been previously shown to be enriched in genes expressed within the central nervous system (CNS) and, more specifically, neuronal cell types^14,17^, and was therefore included as a measure of robustness.

**Figure 1.**
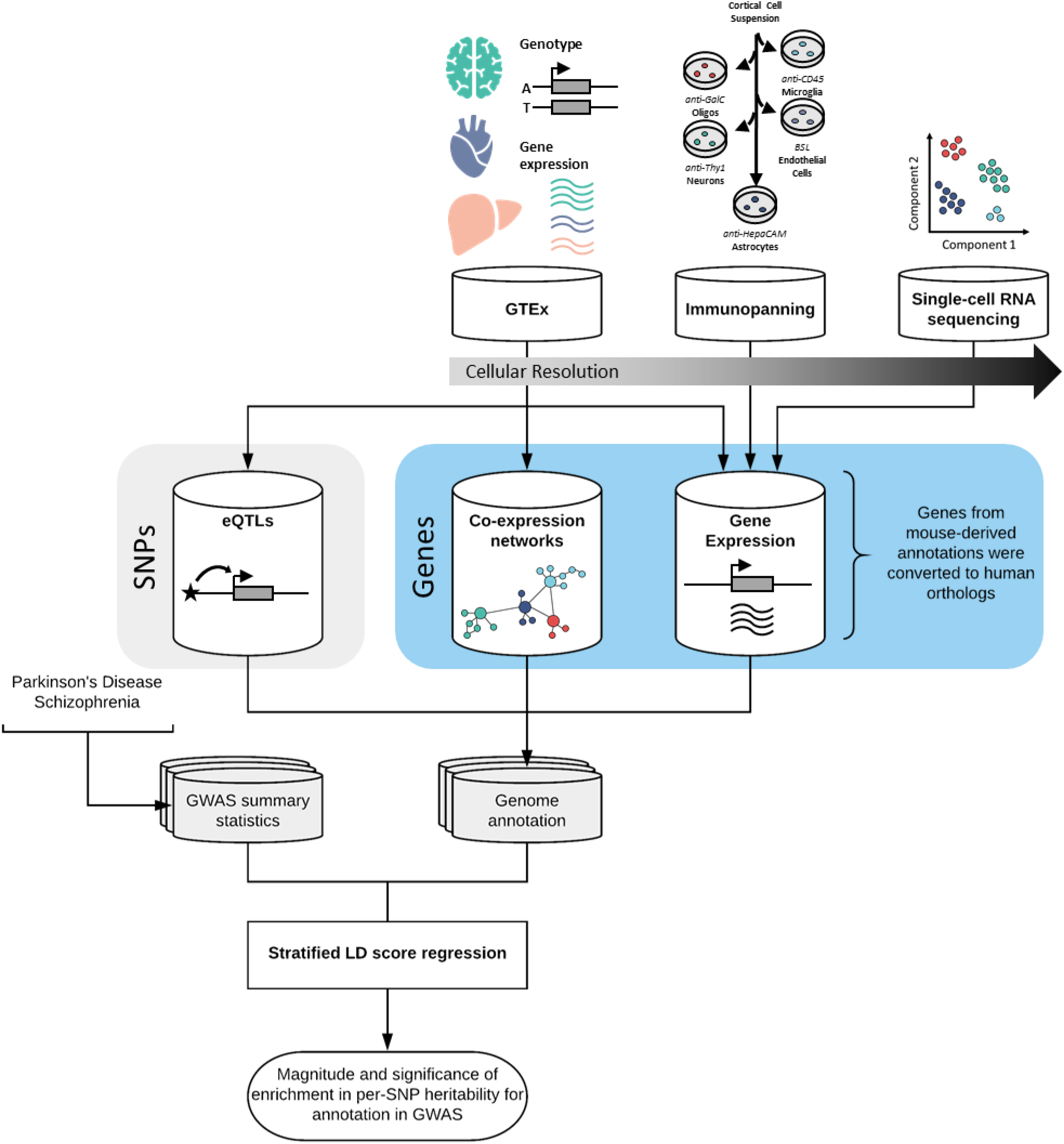
Overview of approach and datasets used. This study compiled several brain-related genomic annotations reflecting tissue- and cell-type-specific activity, using data generated by the GTEx project^19^, the Barres group^20^ and the Linnarsson group^21^. These annotations, each of which varied in their cellular resolution, included: tissue-specific eQTLs (reflecting the effect of genetic variation on gene expression); tissue-specific co-expression networks (reflecting the connectivity of a gene to all other expressed genes in the tissue), and tissue- and cell-type-specific gene expression. All annotations were constructed in a binary format (1 if the SNP is present within the annotation and 0 if not). For annotations where the primary input was a gene, all SNPs with a minor allele frequency > 5% within ± 100kb of the transcription start and end site were assigned a value of 1. For more details of how each individual annotation was generated see Methods. Stratified LDSC was then used to test whether an annotation was significantly enriched for the common-SNP heritability of PD and SCZ.

### PD heritability does not enrich in genomic regions specifically expressed or regulated in human brain

It is well recognised that regional differences in gene expression within human brain and related co-expression modules are driven by differences in the type and density of specific cell types^18^. Therefore, we first used regional data as a means of capturing major cellular profiles. This information, comprising sets of human tissue-specific genes generated by Finucane *et al.* with GTEx gene expression^17,19^, had the advantage of being comprehensive in terms of sampling across the human CNS, and being robust in that greater than 63 independent samples contributed to the generation of each profile. We confirmed that SCZ heritability was significantly enriched in all 13 brain regions relative to all other tissues, as previously demonstrated by Finucane *et al.*^17^ using the 2014 SCZ GWAS (Figure 2A, Supplementary Table 1). In contrast, no tissues were enriched for PD heritability, although spinal cord and substantia nigra approached the Bonferroni significance threshold (threshold p-value = 4.72 × 10^-4^; spinal cord (cervical c-1), p-value = 3.53 × 10^-3^; substantia nigra, p-value = 8.96 × 10^-4^). Our comparison of PD and SCZ GWAS iterations across the years revealed the robust nature of the CNS enrichment in SCZ, which was apparent in the first and smallest SCZ GWAS (Supplementary Figure 1). Furthermore, increasing GWAS sample sizes was associated with co-efficient p-values becoming more significant, particularly for CNS-related tissues. Interestingly, we also observed an ordering of tissues, with brain regions of greater relevance to disease pathology demonstrating the most significant co-efficient p-values in the largest GWAS iterations (e.g. substantia nigra in PD and frontal cortex in SCZ).

**Figure 2.**
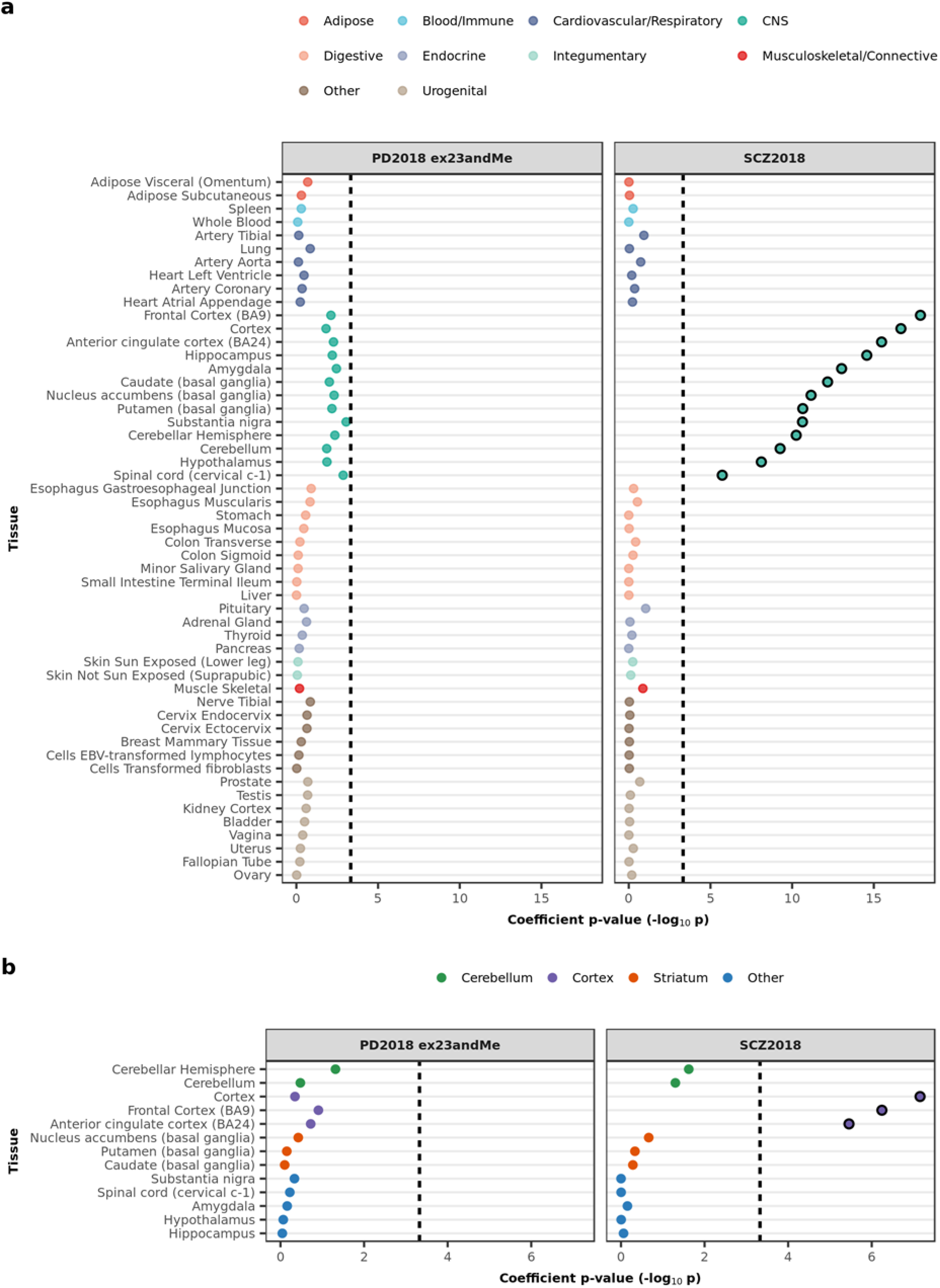
Enrichment of PD and SCZ common-SNP heritability in tissue-specific gene expression annotations as used in Finucane *et al.*^17^. **A**) Stratified LDSC analyses showed a significant enrichment of SCZ heritability in all GTEx brain regions but no enrichment of PD heritability. GTEx tissue annotations represent the top 10% most upregulated genes in each tissue with respect to the remaining tissues, excluding those from a similar tissue category. **B**) Stratified LDSC analyses showed a significant enrichment of SCZ heritability in cortical brain regions, but no enrichment for PD heritability. GTEx brain-only annotations represent the top 10% most upregulated genes in each brain region with respect to the remaining regions. The black dashed lines indicate the cut-off for Bonferroni significance (**A**, p < 0.05/(2 × 53); **B**, p < 0.05/(2 × 13)). Bonferroni-significant results are marked with black borders. Results for previous iterations of the PD and SCZ GWASs are displayed in Supplementary Figure 1 and 2. Numerical results are reported in Supplementary Table 1.

However, due to the way these annotations were constructed, related tissues (e.g. brain regions) have overlapping gene sets and therefore may appear enriched as a group. To differentiate among brain regions, we used fine-scale brain expression data generated by Finucane *et al.* from a brain-only analysis of the 13 GTEx brain regions^17^. We confirmed significant enrichments in the cortex relative to other brain regions for SCZ, but saw no enrichments for PD (Figure 2B, Supplementary Table 1).

We also compared the PD and SCZ GWAS results to sets of blood- and brain-specific eQTLs derived from GTEx. We demonstrated an enrichment of SCZ heritability in brain-specific eQTLs and blood-specific eQTLs, but no enrichment of PD heritability in either eQTL annotation (Figure 3A, Supplementary Table 2). A comparison of eQTLs specific to each brain region revealed no preferential enrichment of disease heritability in one region relative to the others (Figure 3B, Supplementary Table 2). In summary, these analyses revealed no enrichment of PD heritability brain annotations, while in contrast, SCZ heritability was highly enriched in both global and specific regional brain annotations.

**Figure 3.**
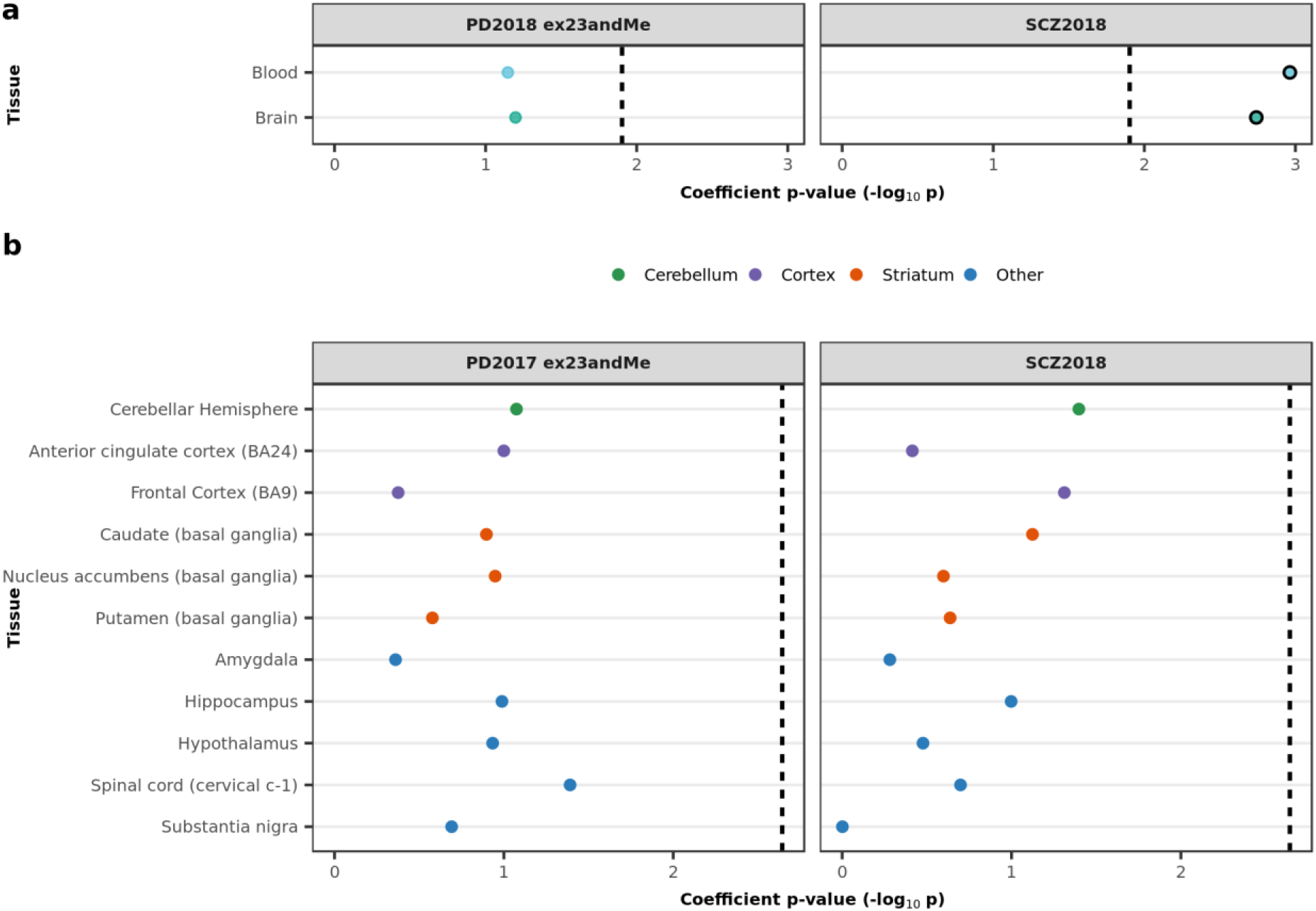
Enrichment of PD and SC common-SNP heritability in tissue-specific eQTL annotations. **A**) Stratified LDSC analyses showed a significant enrichment of SCZ heritability in brain-specific and blood-specific GTEx eQTLs. **B**) A within-brain analysis of GTEx eQTLs showed no significant enrichment of PD and SCZ heritability in one region relative to others. In both analyses, eQTLs were assigned to a tissue/brain region based on their effect size (i.e. the absolute value of the linear regression slope). The black dashed lines indicate the cut-off for Bonferroni significance (**A**, p < 0.05/(2 × 2); **B**, p < 0.05/(2 × 11)). Bonferroni-significant results are marked with black borders. Results for previous iterations of the PD and SCZ GWASs are displayed in Supplementary Figure 3. Numerical results are reported in Supplementary Table 2.

### PD heritability does not enrich in brain-related cell-type-specific annotations

Given the lack of enrichment of PD heritability in global and regional brain annotations, we wondered whether cellular heterogeneity may be masking signals, and provided more cell-type-specific information the enrichment would become more apparent. Thus, to address the relative importance of brain cell types in PD and SCZ, we generated cell-type-specific annotations from three types of brain-related cell-type-specific data: bulk RNA-sequencing from the Barres group of immunopanned cell types from human temporal lobe cortex^20^; single-cell RNA-sequencing from the Linnarsson group of the adolescent mouse nervous system^21^; and finally, cell-type modules inferred from human tissue-level co-expression networks^22^. Genes were assigned to cell types by fold enrichment (i.e. mean expression in one cell type divided by the mean expression in all other cell types) or module membership in the case of co-expression (module membership is a measure of how correlated a gene’s expression is with respect to a module’s eigengene)..

Each of these datasets came with advantages and disadvantages, which motivated our decision to use all three. The Barres data was based on the analysis of human tissue; however, it covered only one brain region, was derived from a small number of individuals (n = 14) who all had an underlying neurological disorder (epilepsy, stroke, glioma), and lacked cellular detail. While the cell-type-specific data provided by the Linnarsson group covered both the central and peripheral nervous system, and contained remarkable cellular detail, it was mouse in origin. Cell-type modules also covered several brain regions, were based on large sample sizes, and importantly, were human in origin. Nevertheless, they were inferred cell types, the definition of which was strongly dependent on the quality of the cell-type markers used to identify them.

Using immunopanning data, we identified a neuronal enrichment for SCZ heritability, but no cell-type enrichment for PD (Figure 4A, Supplementary Table 3). We questioned whether this lack of cell-type enrichment in PD may result from sampling a tissue which is typically affected only in the later stages of sporadic PD^2^. Thus, we analysed a subset of mouse single-cell data representing tissues affected in earlier stages of sporadic PD, including the enteric nervous system, the substantia nigra and the basal ganglia. Once again, we found no cell-type enrichment for PD heritability (Figure 4B, Supplementary Table 3). Conversely, we demonstrated a significant enrichment of SCZ heritability in three types of GABAergic medium spiny neurons (MSNs): MSN2, MSN3 and MSN5. This is consistent with the findings reported by Skene *et al.*^23^. Common to all three types of MSN is that they express the D_2_ dopamine receptor, a common target of antipsychotic drugs used in SCZ therapy^24^.

**Figure 4.**
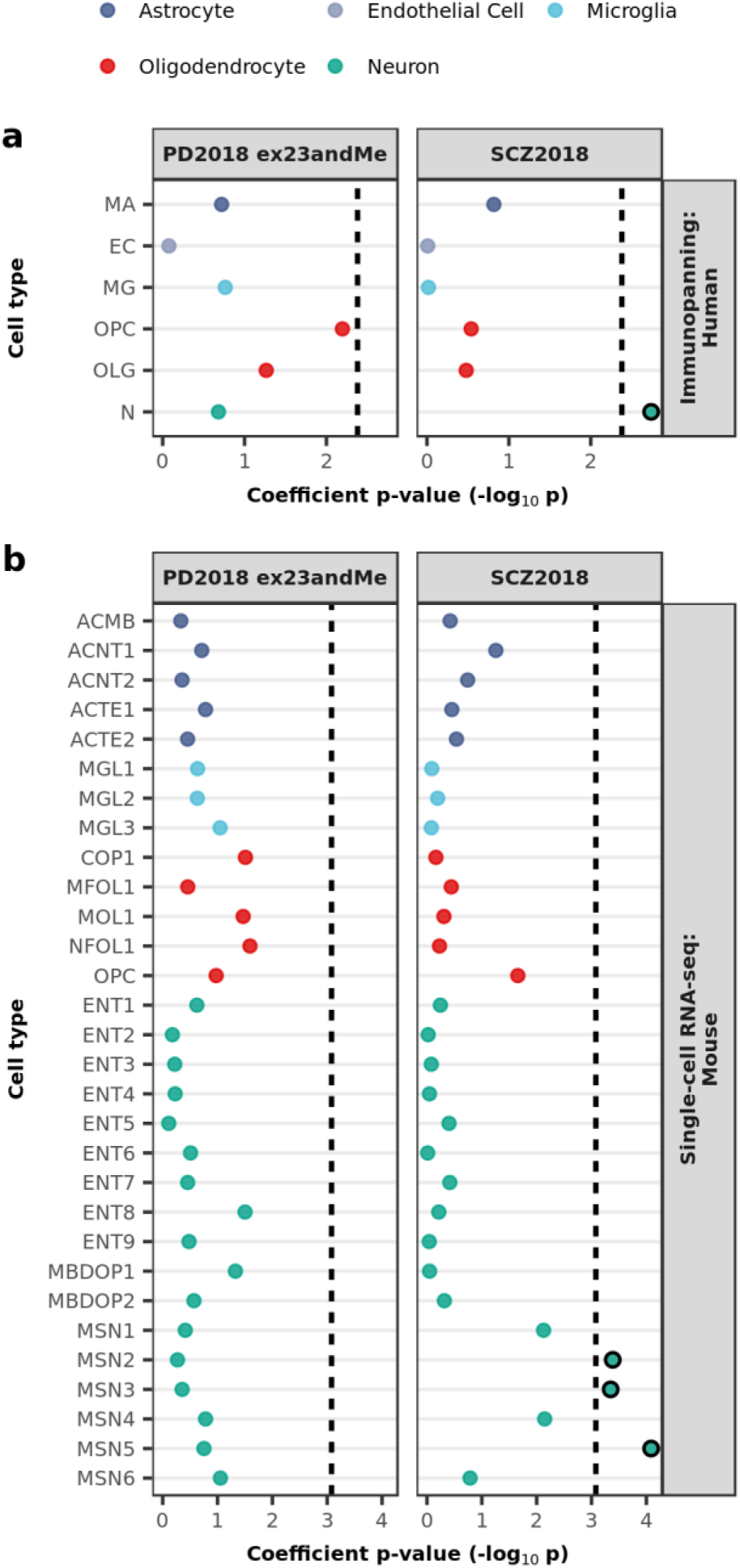
Enrichment of PD and SCZ common-SNP heritability in brain-related cell-type-specific gene expression annotations. Stratified LDSC analyses using cell-type-specific annotations derived from bulk RNA-sequencing of immunopanned cell types from human temporal lobe cortex (**A**) and single-cell RNA-sequencing of the adolescent mouse nervous system (**B**) demonstrated an enrichment of SCZ heritability in neuronal cell types (in particular, medium spiny neurons), but no cell-type enrichment for PD. All cell-type annotations were generated using the top 10% of enriched genes within a cell type compared to all others. Bonferroni significance (**A**, p < 0.05/(2 × 6); **B**, p < 0.05/(2 × 30)). The black dashed lines indicate the cut-off for Bonferroni-significant results are marked with black borders. Results for previous iterations of the PD and SCZ GWASs are displayed in Supplementary Figure 4. Numerical results and cell-type abbreviations are reported in Supplementary Table 3.

To our knowledge, there is currently no single-cell RNA-sequencing data for human striatum or substantia nigra, so we sought to validate our findings using cell-type modules inferred from co-expression networks constructed from human tissue-level expression data of the frontal cortex, putamen and substantia nigra. We observed no significant enrichments for PD heritability in any modules, while SCZ heritability was enriched in several neuronal modules, including: brown and turquoise modules in the frontal cortex; blue and darkmagenta modules in the putamen; and cyan and darkgrey modules in the substantia nigra (Figure 5, Supplementary Table 4). All of these modules were enriched for markers of pyramidal S1 neurons, which have previously been associated with SCZ^23^. Furthermore, some modules (brown, turquoise, darkmagenta and cyan) were enriched for markers of interneurons and dopaminergic neurons, both of which are implicated in SCZ^23,24^.

**Figure 5.**
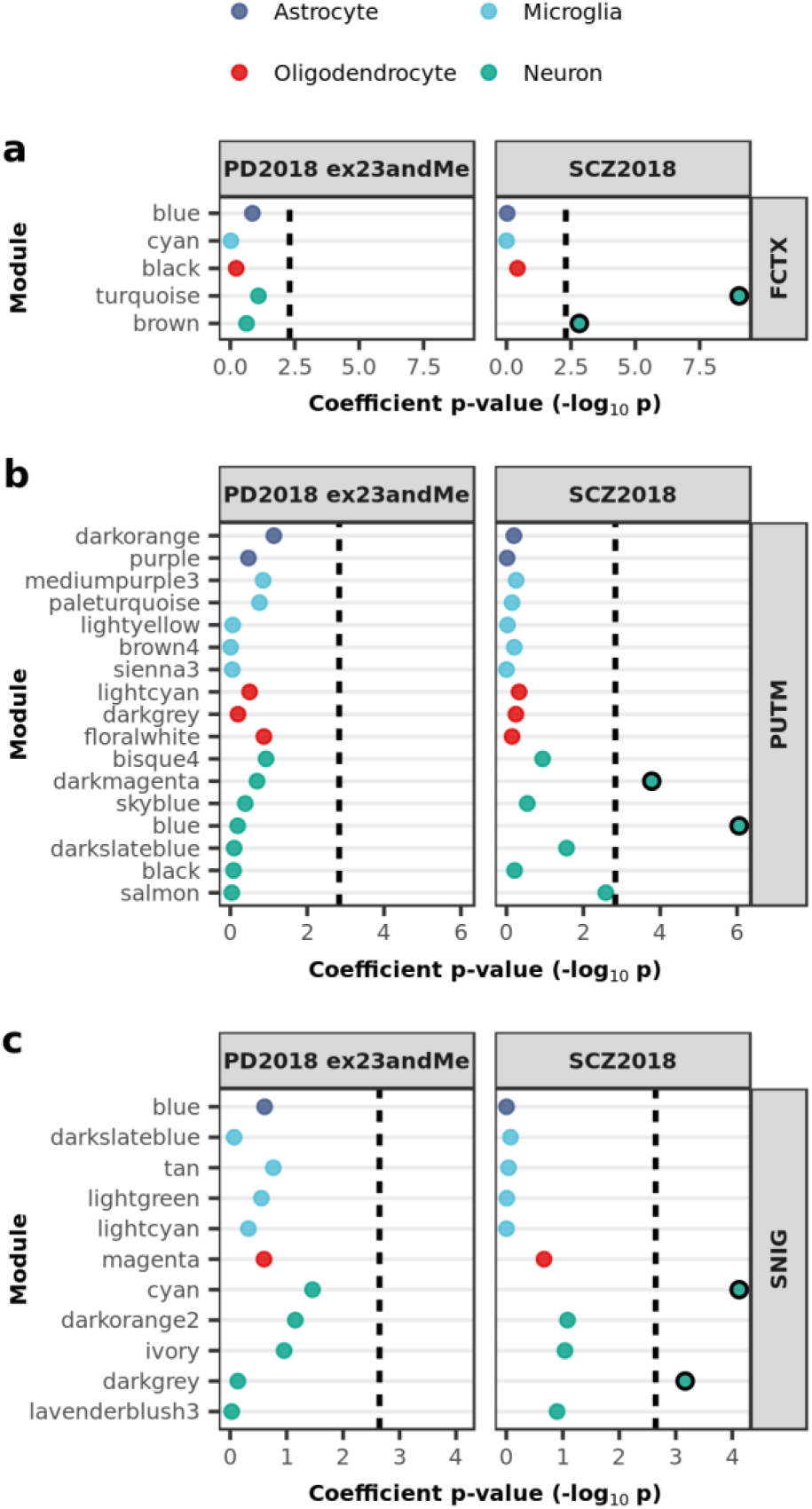
Enrichment of PD and SCZ common-SNP heritability in cell-type modules inferred from human tissue-level co-expression networks. Stratified LDSC analyses using cell-type-specific co-expression modules from frontal cortex (**A**), putamen (**B**), and substantia nigra (**C**) demonstrated a significant enrichment of SCZ heritability in certain neuronal modules across all three tissues, but no enrichment for PD heritability. Genes were assigned to cell-type modules by module membership. The black dashed lines indicate the cut-off for Bonferroni significance (**A**, p < 0.05/(2 × 5); **B**, p < 0.05/(2 × 17); **C**, p < 0.05/(2 x 11)). Bonferroni-significant results are marked with black borders. Results for previous iterations of the PD and SCZ GWASs are displayed in Supplementary Figure 5. Numerical results and module descriptions are reported in Supplementary Table 4. FCTX, frontal cortex; PUTM, putamen; SNIG, substantia nigra.

### PD susceptibility genes do not enrich in brain-related cell types

To ensure rigour, we attempted to identify cell types of importance to PD in a separate analysis using expression-weighted cell-type enrichment (EWCE). This method statistically evaluates whether a set of genes has higher expression in one cell type than expected by chance. Using the same subset of clusters from the Linnarsson single-cell RNA-sequencing, cell-type specificity values were computed for each gene (i.e. proportion of expression for a gene in a given cell type), and cell-type enrichments of PD susceptibility genes implicated by common-variant studies were estimated (Figure 6A, see Methods). We found no significant enrichment of PD susceptibility genes in any of the major cell-type classes (Figure 6B, Supplementary Table 5) or their cell subtypes (Figure 6C, Supplementary Table 5).

**Figure 6.**
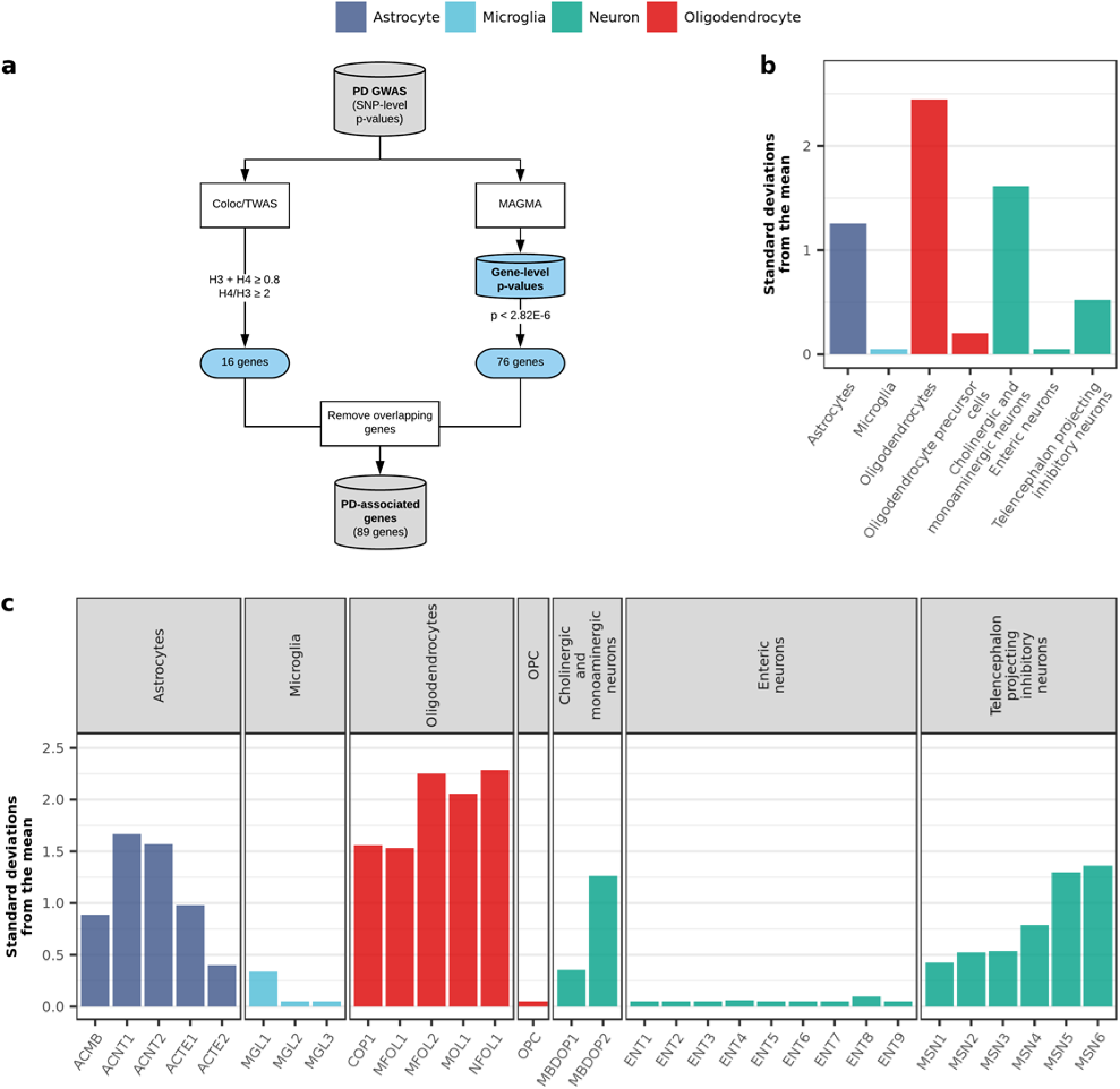
PD susceptibility genes do not enrich in brain-related cell types. **A**) PD susceptibility genes were derived from MAGMA analyses and a study attempting to prioritise genes in PD using TWAS and colocalisation analyses^41^. Genes overlapping between the two sets were removed, resulting in a list of 89 genes. Bootstrapping tests performed using the EWCE method revealed no enrichment of PD susceptibility genes in the major cell-type classes (**B**) or their cell subtypes (**C**) from the Linnarsson single-cell RNA-sequencing dataset. Gene lists and numerical results are available in Supplementary Table 5.

In summary, our EWCE and stratified LDSC analyses would suggest that PD heritability/susceptibility cannot be attributed to a specific cell type (amongst those tested), unlike what has been observed by us and others for SCZ^23^, wherein a limited set of neuronal cell types have been implicated.

### PD heritability enriches in lysosomal genes sets which are specifically expressed in astrocytic and microglial cell subtypes

Risk loci can operate in several manners, including: a cell-type-/tissue-specific manner, which is only detectable if measured in the “correct” cell type/tissue, or in a pathway-specific manner, which one might expect to be detectable across more than one cell type/tissue. Given our inability to implicate a cell type in PD, we wondered whether the latter scenario of pathway-specific risk might be applicable in PD.

To address this question, we applied stratified LDSC to gene sets implicated in PD by Mendelian forms of PD, functional assays performed in the context of PD-associated mutations, such as the A53T missense mutation in *SNCA*, and rare-variant studies of sporadic PD^25–30^. In particular, we focused on gene sets associated with autophagy^26,27^, the lysosomal system^28^ and mitochondrial function^29,30^. Our gene sets were derived either from Gene Ontology terms (autophagy) or curated gene databases (lysosomal, hLGDB; mitochondrial, MitoCarta 2.0; see Methods), developed using literature curation (with a focus on unbiased proteomic studies) and experimental approaches. The overlap between these gene sets was relatively low (Supplementary Figure 6). We identified a significant enrichment of PD heritability in the lysosomal gene set (Figure 7A, Supplementary Table 6). As with previous stratified LDSC analyses, we included SCZ for comparison purposes and found no enrichment of SCZ heritability in any of the assessed gene sets.

**Figure 7.**
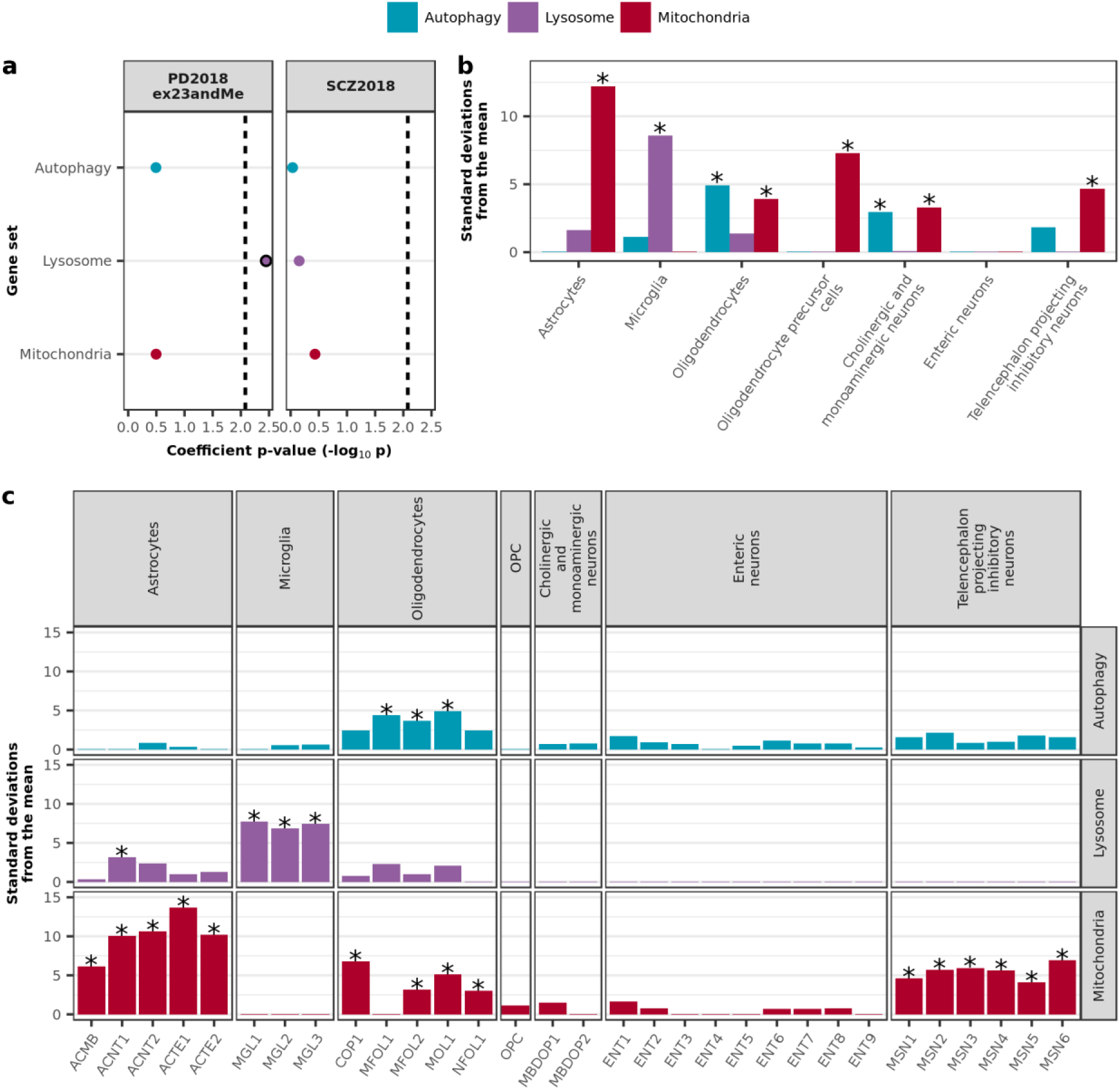
PD heritability enriches in lysosomal gene sets which are specifically expressed in astrocytic and microglial cell subtypes. **A**) Stratified LDSC analyses using gene sets implicated in PD demonstrated a significant enrichment of PD heritability in the lysosomal gene set. The black dashed lines indicate the cut-off for Bonferroni significance (p < 0.05/(2 × 3)). Bonferroni-significant results are marked with black borders. Bootstrapping tests performed using the EWCE method demonstrated enrichment of autophagy, lysosomal and mitochondrial gene sets in specific cell-type classes (**B**) and their cell subtypes (**C**) from the Linnarsson single-cell RNA-sequencing dataset. Asterisks denote significance at p < 0.05 after correcting for multiple testing with the Benjamini-Hochberg method over all gene sets and cell types tested. Gene lists and numerical results are reported in Supplementary Table 6.

Using the same gene sets together with EWCE, we also evaluated whether these PD-implicated gene sets were specifically expressed in any of the Linnarsson cell-type classes and their cell subtypes. Autophagy and lysosomal gene sets were significantly enriched in a limited number of major cell-type classes, with autophagy enriched in oligodendrocytes and cholinergic/monoaminergic neurons, and lysosomal enriched in microglia (Figure 7B, Supplementary Table 6). The mitochondrial gene set, on the other hand, was significantly enriched in almost all cell-type classes, including astrocytes, oligodendrocytes, oligodendrocyte precursor cells, cholinergic/monoaminergic neurons and telencephalon projecting inhibitory neurons. As expected, analyses performed on cell subtypes predominantly reflected that performed on the overarching cell-type classes, with significant pathway enrichments observed in cell subtypes associated with the pathway-enriched cell-type classes (Figure 7C, Supplementary Table 6). For example, all three microglial subtypes (MGL1-3, representing one baseline and two activated microglial subtypes), were enriched for lysosomal genes. Interestingly, the subtype analyses revealed a significant enrichment of the lysosomal gene set in one astrocytic subtype, ACNT1 (non-telencephalon astrocytes, protoplasmic), which was not reflected when using the major cell-type classes, suggesting there may be some pathway specificity within cellular subtypes.

Taken together these findings provide support for the view that in contrast to the genetic structure of SCZ, PD risk loci operate in a more global manner, with effects on a range of cell types.

## Discussion

One of the most striking features of PD is the specificity of its neuropathology and clinical symptoms, which has implicated α-synuclein biology in dopaminergic neurons of the substantia nigra pars compacta as a key component of the disease^1,2^. This stands in stark contrast to SCZ, which has a very heterogeneous clinical phenotype and lacks a characteristic neuropathology^31,32^, with a notable absence of pathological lesions and no reported overall neuronal loss^33^. The apparent cellular specificity of PD has encouraged researchers to hypothesise that selective vulnerability is prompted by the action of risk loci in specific cell types; in other words, it is the nature of the cell type itself, which renders it vulnerable. However, given the interrelated nature of brain regions, apparently specific and reproducible patterns of abnormality could also be the result of a more global effect that exposes functional systems (e.g. neural networks) at different times along a disease’s natural history, a view now put forward by several independent groups^34–36^. That is, risk loci may not necessarily lie in cellular subtypes or individual brain regions, but in global cellular processes to which cellular subtypes have varying vulnerability.

Addressing the question of cellular specificity in sporadic PD in a meaningful manner is now possible due to increasing GWAS sample sizes, increased availability of cell-type-specific gene expression data, and the recent development of robust methodologies. In this study, we used stratified LDSC and EWCE together with several brain-related genomic annotations to connect common-variant genetic findings for PD to specific brain cell types, with SCZ included for comparison purposes. We show that PD heritability does not enrich in global brain annotations or brain-related cell-type-specific annotations, as one might expect if cellular heterogeneity was masking the signal. In contrast, SCZ heritability significantly enriches in global and regional brain annotations and in select neuronal cell types, in line with previous results^17,23^.

One might argue that the lack of PD heritability enrichment in any cell-type-specific categories could be due to PD having a relatively low estimated total heritability; PD heritability estimates range between 20-27% ^9,37,38^. However, we suggest that this is not a complete explanation as significant enrichments have been observed in other GWASs with relatively low overall heritability estimates. For example, in the original stratified LDSC paper they observe enrichment of genomic overlap of histone modifications for the CNS in the ever-smoked GWAS, specifically in the inferior temporal lobe, and they observed enrichment of fetal brain regulatory features for age at menarche^39^.

Considering our inability to attribute PD heritability/susceptibility to a specific brain-related cell type, we also applied stratified LDSC and EWCE to gene sets implicated in PD (autophagy, lysosomal and mitochondrial gene sets), all of which can be considered global pathways. Here we show a significant enrichment of PD heritability in a lysosomal gene set, which is specifically expressed in astrocytic and microglial subtypes, providing support for the view that PD is a disorder of global pathways working across various cell types, as opposed to specific cell types themselves driving disease risk.

With these results in mind, it is tempting to speculate that PD presents genetically as more of a systemic disorder, with a bias to brain pathology, as opposed to a primary brain disorder. In support of this view, PD-associated risk variants have been found associated with monocytes and the innate immune system^13,40,41^, in addition to lymphocytes, mesendoderm, liver- and fat-cells^42^. Recent work has also demonstrated a causal relationship between BMI and PD^43^, which together with the re-purposing of exenatide (a glucagon-like peptide-1 receptor agonist currently licensed for the treatment of type 2 diabetes) for the potential treatment of PD^44^, highlights the need to look beyond the brain and selective neuronal vulnerability.

There are several caveats to this study, namely the quality of our annotations, the strategies employed to generate them and, perhaps most critically, the annotations we cannot account for. First, the quality of our annotations is especially pertinent in the case of the gene sets used to reflect various PD-implicated pathways. While lysosomal and mitochondrial gene sets were derived from rigorously curated gene databases, with a focus on unbiased proteomic and localisation studies, the autophagy list stemmed from Gene Ontology, which has not undergone the same meticulous curation. The noise introduced by potentially inaccurate annotation could affect our ability to detect heritability enrichments.

Second, our strategy for creating cell-type-specific profiles primarily involved gene expression data and assumed disease relevance only if disease heritability enriched for SNPs within genes with high specific expression. This approach together with the use of GTEx eQTLs will likely capture regulatory SNPs in close proximity to genes of interest. However, as demonstrated in a recent study from Hormozdiari *et al.* how one chooses to construct an eQTL annotation is fraught with challenges^45^ and we recognise that our approach may have produced conservative enrichment estimates. Perhaps more importantly though, our strategy for creating cell-type-specific profiles does not account for the effect of regulatory SNPs that function at longer distances to impact upon gene expression. At present, our ability to address this issue is limited since detecting trans-acting eQTLs has proven to be challenging^46^, especially in human brain.

Third, our approach accounts only for cell type and pathway and, moreover, builds on the assumption that cellular diversity can be sufficiently described by discrete cells classes, which a recent single-cell RNA-sequencing study of the hippocampal CA1 area has called into question^47^. In this study, it was suggested that characterisation of cells requires continuous modes of variation in addition to discrete cell classes; that is, some cell classes exist on a common genetic continuum. Inherent within this spectrum is cellular state, which reflects the physiological condition of a given cell, whether it be the degree of differentiation or activation in response to a stimulus. There may be cellular states that we have not assayed or captured which harbour PD heritability enrichments. Furthermore, one would expect preferential enrichment of pathways in specific cell types/subtypes to vary dependent on their physiological profile. In view of increasing evidence for the association between PD and the innate immune system^13,40,41^, we think that cellular state is likely to be an important factor, which we cannot fully assess at this stage.

In conclusion, our results add to a growing body of evidence in support of the view that PD risk loci may not lie entirely in those cell types that display the disease’s characteristic neuropathology, but instead in global cellular processes to which cellular subtypes may have varying vulnerability. This view has significant implications for disease modelling, with a choice of model perhaps based upon the cell type, which best reflects the process of interest, as opposed to the cell type which demonstrates the highest burden of α-synuclein aggregates. Likewise, viewing PD as a systemic disorder may have implications for potential drug re-purposing, as in the case of exenatide. Thus, our work here may have wider implications in terms of understanding neurodegenerative disorders more generally as disorders of key cellular processes rather than disorders driven solely by specific cell types.

## Methods

### Stratified LD score regression (stratified LDSC): assessing the heritability of categories of SNPs

We applied stratified LDSC^39^ (see URLs) to determine if various categories of genomic annotations (marking tissue- or cell-type-specific activity, as summarised in **Annotation datasets**) were enriched for heritability of various GWASs (see **GWAS datasets** below). LDSC exploits the expected relationships between true association signals and surrounding local linkage disequilibrium (LD) to correct out confounding biases, such as cryptic relatedness and population stratification, and arrive at unbiased estimates of genetic heritability within a given set of SNPs (here stratified according to whether they were located within genomic annotation regions). Following the procedure employed by Finucane et al.^39^, we added annotation categories individually to the baseline model (version 1.1, see URLs). We used HapMap Project Phase 3 (HapMap3)^48^ SNPs for the regression, and 1000 Genomes Project^49^ Phase 3 European population SNPs for the LD reference panel. We only partitioned the heritability of SNPs with minor allele frequency >5%, and we excluded the MHC region from analysis due to the complex and long-range LD patterns in this region. To map SNPs to genes, we used the SNPlocs.Hsapiens.dbSNP144.GRCh37 R package (dbSNP build 144 and GRCh37 coordinates)^50^.

For all stratified LDSC analyses, we report a one-tailed p-value (coefficient p-value) based on the coefficient *z*-score outputted by stratified LDSC. A one-tailed test was used as we were only interested in annotation categories with a significantly positive regression coefficient (i.e. the annotation positively contributed to trait heritability, conditional upon the baseline model, which accounts for the underlying genetic architecture). We looked at three versions of PD GWAS summary statistics and four versions of SCZ, and for each set of analyses we corrected for multiple testing of the GWASs across the number of annotation categories, resulting in Bonferroni significance thresholds for each set of analyses.

### Annotation datasets

#### Tissue-specific gene expression

Annotation files were generated by Finucane et al.^17^, using GTEx V6P gene expression^19^, and obtained from Alkes Price’s group data repository (see URLs). Briefly, for each GTEx tissue, genes were ranked by a computed *t*-statistic reflecting their specific expression within that tissue versus all other tissues, excluding those that were from a similar tissue category (e.g. expression in cortex samples was compared to expression in all other tissues except other brain regions; see Supplementary Table 2 from Finucane *et al.*^17^ for t-statistic tissue categories). The top 10% of expressed genes from each tissue was selected and a 100-kb window was added around their transcribed regions to obtain a tissue-specific gene expression annotation. For the within-brain analysis, tissues were restricted to the 13 brain regions found in GTEx, including: amygdala, anterior cingulate cortex (BA24), caudate, cerebellar hemisphere, cerebellum, cortex, frontal cortex (BA9), hippocampus, hypothalamus, nucleus accumbens, putamen, spinal cord (cervical c-1), substantia nigra.

#### Tissue-specific eQTLs

From the GTEx Portal (V7, accessed 04/16/18, see URLs), we downloaded all SNP-gene (expression quantitative trait loci, eQTL) association tests (including non-significant tests) for blood (to allow for a blood-brain comparison) and 11 of the 13 available brain regions^19^. To reduce redundancy across the brain regions, we excluded cortex and cerebellum, and instead included frontal cortex, anterior cingulate cortex and cerebellar hemisphere. We performed an FDR correction for each tissue and included all SNP-gene associations that passed FDR < 5% in our downstream analyses. For the blood-brain comparison, eQTLs from all 11 brain regions were combined to form one brain category. eQTLs that replicated across brain regions were collapsed into one entry and allocated an effect size (i.e. the absolute value of the linear regression slope) equal to that of the maximum effect size observed across the brain regions. Finally, eQTLs were assigned to either blood or brain by their effect size. A similar approach was used for the within-brain analysis, where eQTLs were assigned to one of the 11 brain regions based on effect size.

#### Cell-type-specific gene expression

Cell-type-specific annotations were constructed using gene expression data from the Barres group^20^ and the Linnarsson group^21^ (see URLs), which was generated using bulk RNA-sequencing and single-cell RNA-sequencing, respectively. Due to the disparate nature of the RNA-sequencing methods, each dataset was analysed separately. Common to both analyses was the calculation of an enrichment value for each gene in each cell type. Enrichment was calculated as: gene expression in one cell type divided by the average gene expression across all other cell types. We thereafter selected the top 10% of genes enriched within each cell type and added a 100kb window to reflect the approach used by Finucane *et al.*^17^ When using the Barres data, we averaged gene expression across samples of the same cell type, filtered genes on the basis of an FPKM ≥ 1 in at least one cell type (this equates to ~66% of all genes with FPKM > 0.1, which was set by Zhang *et al.* ^20^ as the threshold for minimum gene expression), and then calculated gene enrichment. Our detection threshold of FPKM ≥ 1 was employed on the basis that smaller thresholds tend to produce large and misleading enrichments^51^. The Linnarsson data was available with gene expression aggregated by sub-cell type/cluster. Genes were filtered on the basis of expression > 0, enrichment was calculated and a subset of the 265 identified clusters were used as annotations. Mouse genes were converted to human orthologs using Biomart.

#### Cell-type-specific co-expression modules

Co-expression networks for frontal cortex, putamen and substantia nigra were constructed using GTEx V6 gene expression^19^, the WGCNA R package^52^ and post-processing with k-means^53^ (see URLs), as described by Botia *et al.*^22^ Modules were assigned to cell types using the userListEnrichment R function implemented in the WGCNA R package, which measures enrichment between module-assigned genes and defined brain-related lists^18,54–57^ using a hypergeometric test. Genes assigned to modules significantly enriched for brain-related cell-type markers of predominantly one cell type with a module membership of ≥ 0.5 were allocated a cell-type “label” of neuron, microglia, astrocyte or oligodendrocyte and considered cell-type specific. Module membership values range between 0 and 1, with 1 indicating that a gene’s expression is highly correlated with the module eigengene. An eigengene is defined as the first principal component of a given module and can be considered representative of the gene expression profiles within the module, as it summarises the largest amount of variance in expression.

#### GWAS datasets

**Table 1.** Summary of GWAS datasets.

| Disease | Author, Year | N cases | N controls | PMID | Reference |
| --- | --- | --- | --- | --- | --- |
| PD | IPDGC Consortium, 2011 | 5,333 | 12,019 | 21738488 | <sup>58</sup> |
| PD | Nalls, 2014 | 13,708 | 95,282 | 25064009 | <sup>59</sup> |
| PD | Nalls, 2018 (excluding 23andMe contributions)<sup>1</sup> | 33,674<br>(18,618 proxy cases from UK Biobank) | 449,037 |  | <sup>15</sup> |
| SCZ | SCZ Consortium, 2011 | 9,394 | 12,462 | 21926974 | <sup>60</sup> |
| SCZ | Ripke, 2013 | 14,395 | 18,705 | 23974872 | <sup>61</sup> |
| SCZ | SCZ Consortium, 2014 (EUR subset) | 33,640 | 43,456 | 25056061 | <sup>62</sup> |
| SCZ | Pardiñas, 2018 | 40,675 | 64,643 | 29483656 | <sup>16</sup> |

<sup>1</sup> Access to PD 2018 summary statistics (excluding 23andMe contributions) was provided by Mike A Nalls, with permissions from IPDGC and SGPD.

#### MAGMA: assessing gene-level enrichment

Gene-level p-values were calculated using MAGMA v1.06 (see URLs)^63^, which tests the joint association of all SNPs in a gene with the phenotype while accounting for LD between SNPs. SNPs were mapped to genes using NCBI definitions (GRCh37, annotation release 105); only genes in which at least one SNP mapped were included in downstream analyses. Gene boundaries were defined as the region from transcription start site to transcription stop site. In addition, we added a window of 35 kb upstream and 10 kb downstream of each gene, as most transcriptional regulatory elements fall within this interval^64^. Furthermore, the MHC region on chromosome 6 (chr6: 25500000 – 33500000, human genome assembly GRCh37) was excluded. The gene p-value was computed based on the mean association statistic of SNPs within a gene, with genome-wide significance set to p < 2.82 × 10^-6^, and LD was estimated from the European subset of 1000 Genomes Phase 3^49^.

### Expression-weighted cell-type enrichment (EWCE): evaluating enrichment of PD-associated genes and gene sets

EWCE (see URLs)^65^ was used to determine whether PD-associated genes or gene sets have higher expression within a particular cell type than expected by chance. As our input, we used the same subset of clusters from the Linnarsson single-cell RNA-sequencing dataset used in stratified LDSC, in addition to a target gene list. For each gene in the Linnarsson dataset, we estimated its cell-type specificity i.e. the proportion of total expression of a gene found in one cell type compared to all cell types, using the ‘generate.celltype.data’ function of the EWCE package. EWCE with the target list was run with 100,000 bootstrap lists. We controlled for transcript length and GC-content biases by selecting bootstrap lists with comparable properties to the target list. P-values were corrected for multiple testing using the Benjamini-Hochberg method over all cell types and gene lists tested. We performed the analysis with major cell-type classes (e.g. “astrocyte”, “microglia”, “enteric neurons”, etc.) and subtypes of these classes (e.g. ACNT1 [“Non-telencephalon astrocytes, protoplasmic”], ACNT2 [“Non-telencephalon astrocytes, fibrous”], etc.). Data are displayed as standard deviations from the mean, and any values < 0, which reflect a depletion of expression, are displayed as 0.

#### PD susceptibility genes

PD susceptibility genes were derived from our own MAGMA analyses and a study attempting to prioritise genes in PD using TWAS and colocalisation analyses (Supplementary Table 1 in ref.^41^). The genes comprising these lists are available in Supplementary Table 5. In the case of MAGMA, only those genes passing genome-wide significance (p < 2.82 × 10^-6^) were used. In the case of TWAS/coloc, only those eQTL-gene associations found within dorsolateral prefrontal cortex tissue, which were both TWAS and coloc hits (as defined in ref.^41^) were used.

#### Gene sets associated with PD

We investigated three gene sets with previous biological support for involvement in PD: autophagy, lysosomal and mitochondrial^25–30^. The autophagy gene set included all genes associated with the Gene Ontology terms: GO:0006914 (“autophagy”) and GO:0005776 (“autophagosome”), as derived from the GO C5 collection of the MSigDB database (v5.2). Lysosomal genes were downloaded from the Human Lysosome Gene Database (hLGDB, see URLs)^66^. All genes reported lysosomal by any of the listed sources (9 of the 16 were unbiased proteomic studies) were used. Mitochondrial genes were obtained from Human MitoCarta 2.0, an inventory of human genes with strong support of mitochondrial localisation based on literature curation, proteomic analyses and epitope tagging/microscopy (see URLs)^67^. The genes comprising these lists are available in Supplementary Table 6. Overlap between gene sets was determined using Intervene, a command line tool and web application that computes and visualises intersections of gene sets (see URLs)^68^.

### URLs

Barres immunopanning, http://www.brainrnaseq.org/; Baseline LDSC annotations, https://data.broadinstitute.org/alkesgroup/LDSCORE/; Expression-weighted cell-type enrichment (EWCE), https://github.com/NathanSkene/EWCE; Finucane GTEx annotations, https://data.broadinstitute.org/alkesgroup/LDSCORE/LDSC_SEG_ldscores/; GTEx Portal, https://www.gtexportal.org/; Intervene, https://asntech.shinyapps.io/intervene/; LDSC, https://github.com/bulik/ldsc/wiki; Linnarsson single-cell RNA-sequencing, http://mousebrain.org/; MAGMA, https://ctg.cncr.nl/software/magma; MitoCarta, https://www.broadinstitute.org/scientific-community/science/programs/metabolic-disease-program/publications/mitocarta/mitocarta-in-0; SCZ GWAS summary statistics, https://www.med.unc.edu/pgc/results-and-downloads; The Human Lysosome Gene Database, http://lysosome.unipg.it/; WGCNA, https://labs.genetics.ucla.edu/horvath/CoexpressionNetwork/Rpackages/WGCNA/; WGCNA hierarchical clustering with k-means, https://github.com/juanbot/km2gcn

## Acknowledgements

RHR was supported through the award of a Leonard Wolfson Doctoral Training Fellowship in Neurodegeneration. MAN was supported by a consulting contract between Data Tecnica International and the National Institute on Aging, NIH, Bethesda, MD, USA. JH and MR were supported by the UK Medical Research Council (MRC), with JH supported by a grant (MR/N026004/) and MR through the award of a Tenure-track Clinician Scientist Fellowship (MR/N008324/1). Full consortia acknowledgements are available in the supplemental materials (Text S2).

## Author Contributions

RHR, JH, SAG and MR conceived and designed the study. RHR analysed data, drafted the figures, and together with SAG and MR wrote the first draft manuscript. JB constructed the co-expression networks. MAN, IPDGC and SGPD helped with the use of Parkinson’s disease GWAS data. All authors contributed to critical analysis of the manuscript.

